# Timescales of influenza A/H3N2 antibody dynamics

**DOI:** 10.1101/183111

**Authors:** Adam J. Kucharski, Justin Lessler, Derek A.T. Cummings, Steven Riley

**Affiliations:** Centre for the Mathematical Modelling of Infectious Diseases, London School of Hygiene & Tropical Medicine, UK.; Department of Epidemiology, Johns Hopkins Bloomberg School of Public Health, Baltimore, MD 21205, USA.; Emerging Pathogens Institute, University of Florida, Gainesville, FL 32610, USA.; MRC Centre for Outbreak Analysis and Modelling, Department of Infectious Disease Epidemiology, School of Public Health, Imperial College London, UK.

## Abstract

Human immunity influences the evolution and impact of novel influenza strains. Because individuals are infected with multiple influenza strains during their lifetime and each virus can generate a cross-reactive antibody response, it is challenging to quantify the processes that shape observed immune responses, or to reliably detect recent infection from serological samples. Using a Bayesian model of antibody dynamics at multiple timescales, we explain complex cross-reactive antibody landscapes by inferring participants’ histories of infection with serological data from cross-sectional and longitudinal studies of influenza A/H3N2 in southern China and Vietnam. We show an individual’s influenza antibody profile can be explained by a short-lived, broadly cross-reactive response that decays within a year to leave a smaller long-term response acting against a narrower range of strains. We also demonstrate that accounting for dynamic immune responses can provide a more accurate alternative to traditional definitions seroconversion for the estimation of infection attack rates. Our work provides a general model for explaining mechanisms of influenza immunity acting at multiple timescales based on contemporary serological data, and suggests a two-armed immune response to influenza infection consistent with competitive dynamics between B cell populations. This approach to analysing multiple timescales for antigenic responses could also be applied to other multi-strain pathogens such as dengue and related flaviviruses.

## Introduction

Immunity against influenza A can influence the severity of disease [1, 2], the effectiveness of vaccination strategies [3], and the emergence of novel strains [4, 5]. Understanding the accumulation of immunity and infection has proven challenging because observed human antibody responses – typically measured by haemagglutination inhibition (HI) assays or microneutralisation titres – reflect a combination of past infections to specific strains and the potentially cross-reactive responses generated by these infections [6]. It has been shown that measurement error in HI assays can lead to uncertainty in the estimation of serological status [7] and cross-reactive antibody dynamics can make it difficult to estimate the true extent of influenza infection during an epidemic [2]. Accurate estimation of attack rates is crucial for estimating influenza burden, and hence the design and evaluation of vaccination campaigns [8].

Although there are established techniques for the analysis of single strain immunising pathogens such as measles [9], potential cross-reactivity between different influenza A strains means serological analysis must account for the dynamics of antibody responses across multiple infections [10]. The concept of an antibody landscape has been put forward as one way to represent the immune response developed as a result of a sequence of processes such as infection, antibody boosting, antibody waning and cross reactivity [11]. Previous work has also used cross-sectional data to explore the life course of immunity by explicitly modelling both the processes of infection and immunity [12]. However, such analysis could not examine antibody mechanisms operating at multiple time-scales. In particular, there have been suggestions that influenza infection leads to ‘back-boosting’, generating a broadly cross-reactive response against historical strains [11, 13, 14]. It has also been suggested that influenza responses are influenced by antigenic seniority, with strains seen earlier in life shaping subsequent antibody responses [15]. This is a refinement on the earlier concept of ‘original antigenic sin’, whereby the largest antibody response is maintained against the first infection of a lifetime [16].

To quantify antibody kinetics over time and estimate historical infections with influenza A/H3N2, we used a dynamic model of immune responses that generated expected titres against specific strains [12] by combining infection history – which was specific for each individual – with an antibody response process that was universal across individuals. We assumed that the response included both a short-term and long-term component. The short-term component consisted of a boost in log-titre following infection, which decayed over time, as well as a rise in log-titre as a result of cross-reaction with antigenically variable strains. The long-term response featured a boost in log-titre, which did not decay, and a separate cross-reaction process that led to increased titres against other strains. Titres were also influenced by antigenic seniority, with later infections generating lower levels of homologous boosting than that generated against strains encountered earlier in life (see Methods). Historical strains were assumed to follow a smooth path through a two-dimensional antigenic space over time [17] (Fig. S1). We fitted this model to two publicly available serological datasets in which participants were tested against a panel of A/H3N2 strains. The first contained cross-sectional data for individuals living in Guangdong province in southern China, collected in 2009 [15, 18]; the second included longitudinal data from Ha Nam in Vietnam [19], with sera collected between 2007–2012 [11, 20].

## Results

Using our serological model, we jointly estimated influenza infection history for each study participant, as well as subsequent antibody response processes and assay measurement variability. Although the contributions of short- and long-term processes to antibody responses cannot be robustly estimated from cross-sectional data [12], simulation studies showed that both time scales were identifiable using a simulated dataset similar to that of the Vietnam samples (Figs. S2–3). We therefore included the short-term dynamic antibody processes in the model when fitting longitudinal data, but not when fitting to cross-sectional data. The fitted model could reproduce both cross-sectional and longitudinal observed titres for each participant (Fig. 1), and it was possible to identify specific years with a high probability of infection and the corresponding antibody profile this infection history had generated (Table S1, Figs. S4–5). Using the longitudinal Vietnam data, we could identify specific years in which individuals had a high probability of infection, particularly during the period of testing (Fig. 1A–I). There was more variability in estimates from the cross-sectional China data, although time periods with a high probability of infection could still be identified (Fig. 1J–L).

**Figure 1:**
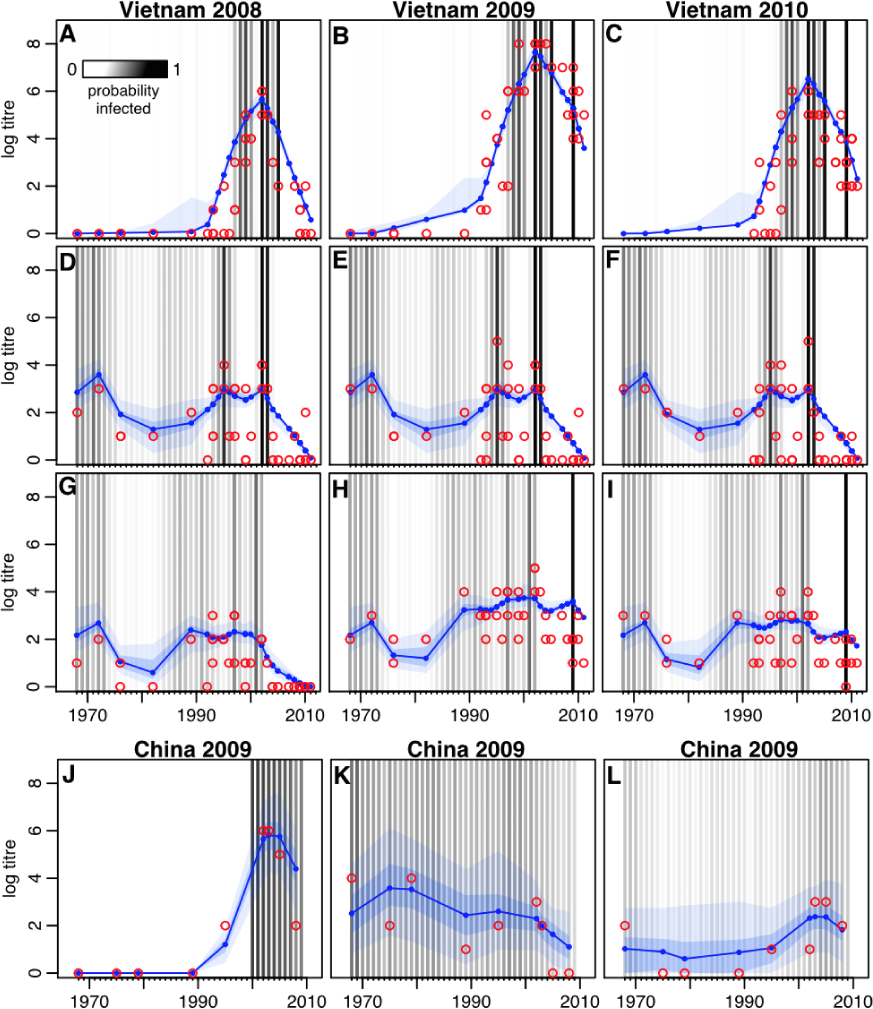
Representative individual-level responses against influenza in Vietnam (A–I) and southern China (J–L). **(A–C)** Strong evidence of infection in 2009, leading to rise in titres and back boost from broad short-term cross-reaction, then decay in following year. Red points show observed titre. Blue lines show median titre in fitted model, with blue regions showing 50% and 95% MCMC credibility intervals. Black lines show samples from the posterior distribution of individual infection histories, with opacity indicating the probability of infection (i.e. proportion of MCMC samples that estimated infection in that year). **(D–F)** No estimated infections between 2008–2010, so titres are at equilibrium. **(G–I)** Infection in 2009 leading to broad boost, with titres generally highest against recent strains **(H)** then decline to equilibrium, with lower mean titres against recent strains as a result of antigenic seniority **(I)**. **(J–L)** Cross-sectional results from southern China, indicating: **(J)** evidence of multiple recent infections; **(K)** decline in titres as a result of antigenic seniority; **(L)** evidence of infections early and late in life.

The model fits to longitudinal data described an antibody response to influenza that is initially dominated by a broadly cross-reactive response, which rapidly decays, leaving a long-term response that cross-reacts only with antigenically similar viruses (Table 1, Figs. S6–7). We estimated that primary infection generated a short-lived boost of an average of 2.69 (95% CrI: 2.50–2.89) units of log-titre against the infecting virus (a fourfold rise would be equivalent to a 2 unit rise in log-titre), and a long-term boost of 1.80 log-titre units (95% CrI: 1.74–1.88). The short-term response decayed quickly: we estimated that the response had reached its final equilibrium level after one year. As the samples were collected at one year intervals, it was not possible to estimate beyond this level of precision, but it suggests that the short-term response contributes to titres on a timescale of less than one year. The timescale of this short-term response is consistent with previous qualitative estimates based on laboratory confirmed infections, which suggested there was a negligible change in titre more than one year post-infection [11, 13, 21].

**Table 1:** Parameter estimates for models fitted to data from southern China and Vietnam. Median estimate shown, with 95% credible intervals in parentheses.

| Parameter | China (2009) | Vietnam (2007–2012) |
| --- | --- | --- |
| Long-term boost ($\mu_1$) | 1.42 (1.12-1.67) | 1.80 (1.74-1.88) |
| Short-term boost ($\mu_2$) | – | 2.69 (2.50-2.89) |
| Long-term cross-reaction ($\sigma_1$) | 0.128 (0.102-0.145) | 0.134 (0.131-0.140) |
| Short-term cross-reaction ($\sigma_2$) | – | 0.0327 (0.0287-0.036) |
| Observation error ($\varepsilon$) | 1.69 (1.55-1.83) | 1.29 (1.27-1.31) |
| Antigenic seniority ($\tau$) | 0.029 (0.021-0.037) | 0.044 (0.039-0.048) |
| Waning ($\omega$) | – | 0.995 (0.973-1.00) |

For the long-term response inferred from longitudinal data, we estimated that crossreactivity between infecting strain and tested strain dropped off at a rate of 0.241 units of log-titre (95% CrI: 0.228–0.261) per unit of antigenic distance between them. The estimated drop was not significantly different from that inferred with the cross-sectional China data: log-titres decreased by 0.182 (0.116-0.237) with each antigenic unit. The broader credible interval for China was largely the result of the coverage of strains tested: 9 strains were tested in the China data, compared with up to 57 in the Vietnam data. For the broader short-term response, the model fitted to longitudinal data suggested crossreactive titres only decreased by 0.088 (95% CrI: 0.074–0.101) with each antigenic unit. This result suggests that short-term titres are influenced by antigenic distance, albeit weakly, and hence provides quantitative support for previous suggestions that the observed broad short-lived boost is part of a memory B cell response [11].

To illustrate the inferred short and long term antibody dynamics against A/H3N2, we used our infection history model to simulate antibody responses following two sequential infections, the first in 1968 and in 1988 (Fig. 2). Following primary infection, individuals would be expected to have raised titres to strains in nearby regions of antigenic space, but these titres would quickly decay to leave a more localised long-term response. Upon secondary infection, a similar boost in titres would be observed, which would not be present in tests conducted in subsequent years. This highlights the importance of accounting for multiple-time scales when analysing immune assay data: in simulations, serology taken in 1988 indicated a rise in titre to the first infecting strain compared to serology between 1969–1987, and showed detectable titres against all strains in the region of antigenic space between the two infecting strains (Fig. 2F). However, serology taken one year later only displayed localised responses against the infecting strains (Fig. 2H). Depending on time of sampling, our results suggest it would be possible to observe either longitudinal increases or decreases in log-titres against previously seen strains or stable log-titres [6].

**Figure 2:**
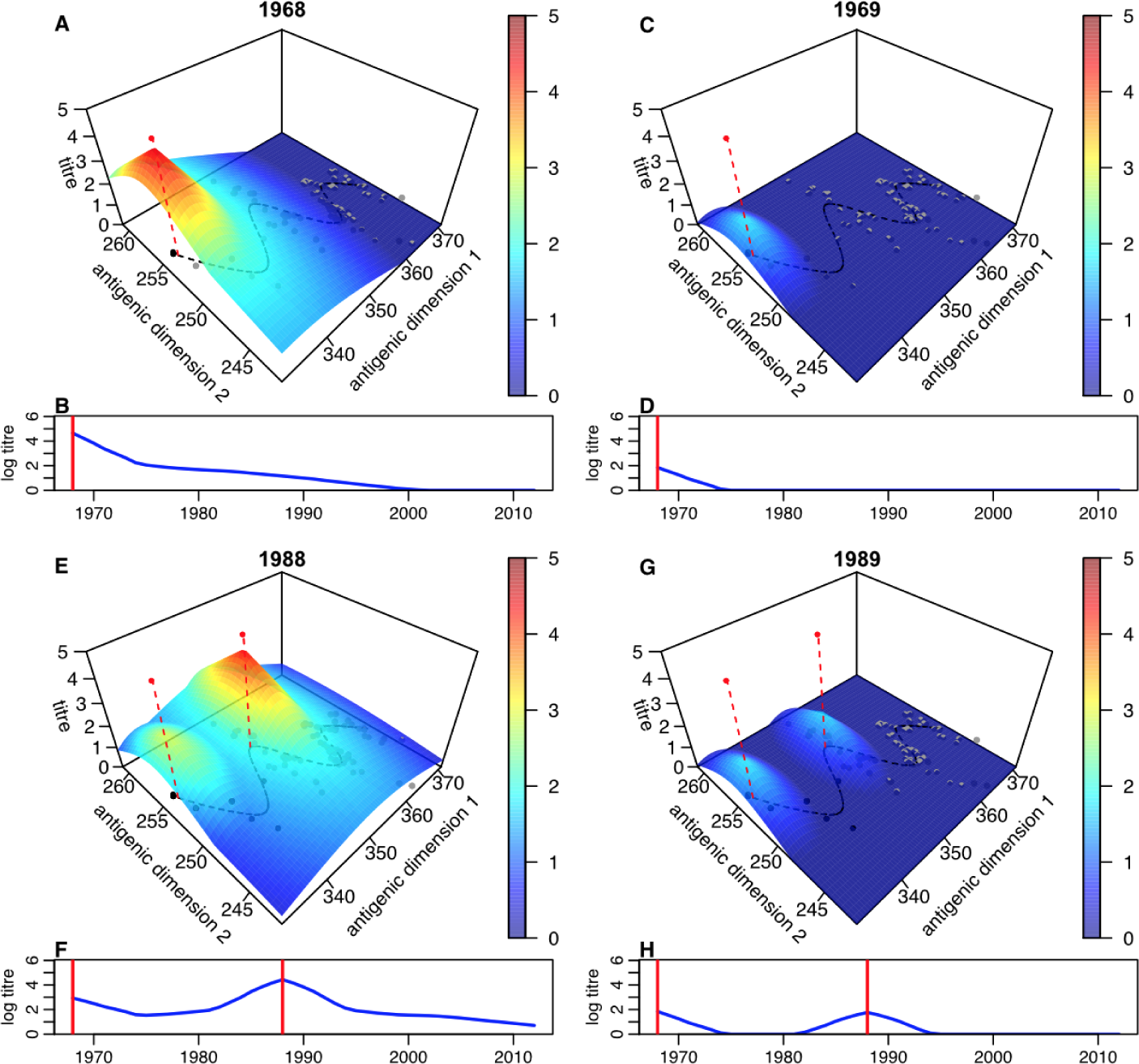
Expected titres against strains at different points in antigenic space for a given infection sequence. **(A)** Simulated log-titres against different strains in antigenic space following a single infection in 1968, with test conducted in 1968. Parameters are drawn from the maximum a posteriori model estimate. Red vertical dashed line shows antigenic location of infecting strain. Black points at the base show location of strains isolated up to this year; grey points show location of strain isolates in subsequent years; black dashed line shows antigenic summary path used to fit model (Figure S1). **(B)** Estimated titres along the antigenic summary path (dashed black line in **(A)**). Red line shows year of infection. **(C)** Simulated log-titres following on single infection in 1968, with test conducted in 1969. **(D)** Estimated titres along antigenic summary path in 1969. **(E)** Simulated log-titres following infections in 1968 and 1988, with test conducted in 1988. **(F)** Estimated titres along antigenic summary path in 1988. **(G)** Simulated log-titres following infections in 1968 and 1988, with test conducted in 1989. **(H)** Estimated titres along antigenic summary path in 1989.

As well as examining antibody dynamics, we reconstructed historical annual attack rates. In simulation studies, the model could accurately recover attack rates from Vietnam like serological data, particularly for recent years (Fig. 3A). Estimates of attack rates based on the traditional gold-standard of a four-fold rise in titre underestimated the actual simulated values (Fig. 3B), and an overestimate was obtained if a two-fold rise in titre was considered instead [7]. This suggests that commonly used metrics could substantially bias estimates of population-level attack rates, and hence conclusions about the potential extent of herd immunity and required vaccination coverage. In contrast, estimates from our joint inference framework consistently recovered the true simulated infection dynamics during the period of sampling (Fig. 3B, inset). Applying our inference framework to real data from Vietnam to estimate annual attack rates (Fig. 3C), we found that estimates were consistent with observed epidemiological dynamics in Vietnam between 2008–2012, as measured by the number of influenza A/H3N2 isolates during the testing period (Fig. 3D). The correlation between model estimates and observed values was *ρ*=0.996 (p<0.001), with a weaker association when a two-fold rise (*ρ*=0.862, p=0.14) or four-fold rise (*ρ*=0.799, p=0.20) was used to estimate attack rates. Most of the uncertainty in attack rate estimates resulted from individuals with multiple estimated infections; there was little variation in estimated number of infections when individuals had fewer than around eight median infections (Fig. S8).

**Figure 3:**
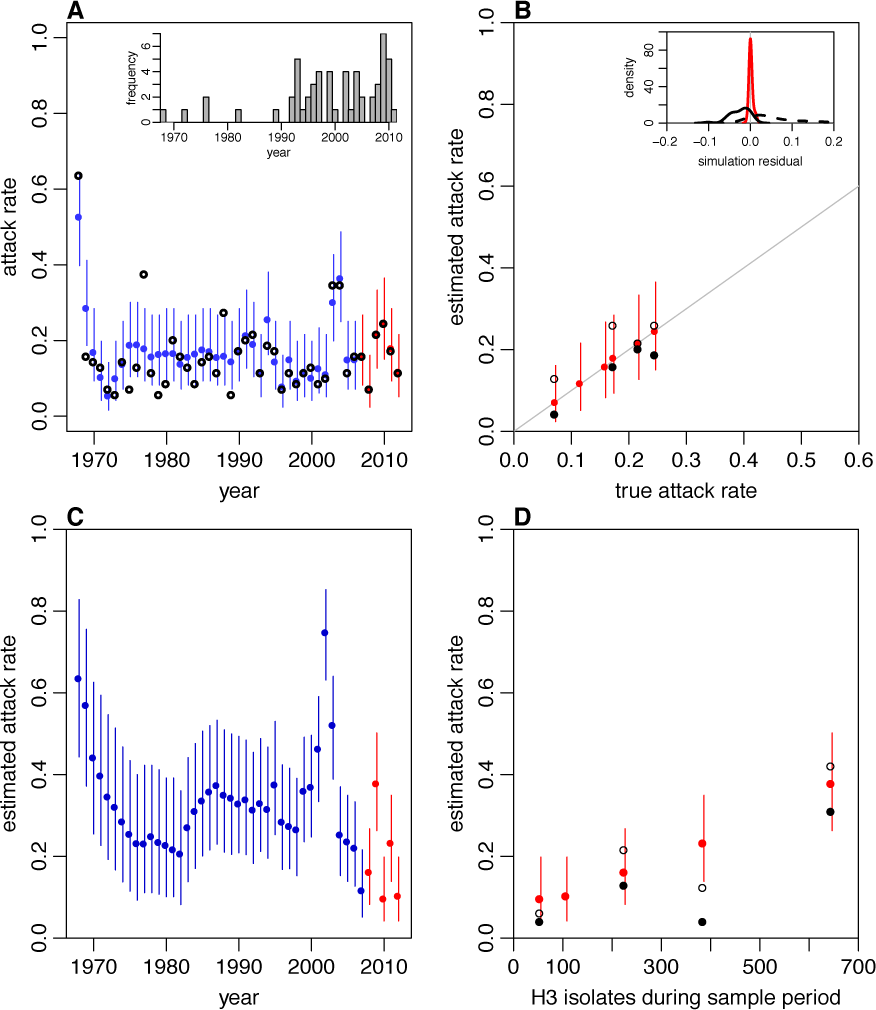
Estimation of influenza A/H3N2 attack rates. **(A)** Inference of attack rates using simulated data for 69 participants, with same strains as tested in HaNam data. Main plot: Blue lines show estimated attack rate with binomial confidence interval; red lines show attack rates in years when samples were taken; black circles show true attack rate in original simulation. Inset: year of circulation for the 57 test strains used, which included repeats in some years. **(B)** Main: accuracy of attack rate estimates in **(A)** was high for years in which serological samples were collected (shown as red dots). Hollow black points show attack rate based on two-fold rise in titre against strain in that year (points shown for years 2008–11, which had sufficient test strains or samples to perform this calculation); solid points show attack rate based on four-fold rise. Inset: distribution of differences between estimated and actual attack rates in same years across ten simulation studies. Red line indicates estimates from model; dashed black line shows estimates based on two-fold rise in titre; solid black line shows estimates based on four-fold rise. **(C)** Proportion of Vietnam study population estimated to have been infected in each year based on real data. Blue lines show estimated attack rate with binomial confidence interval; red lines show attack rates in years when samples were taken. **(D)** Accuracy of attack rates estimates using different methods. Main plot shows model estimates of attack rates in 2008–2012 (red points in **(A)**) and number of positive H3 isolates reported in Vietnam during the same intervals as the samples were taken. Hollow black points show attack rate based on two-fold rise in titre against strain in that year (point not shown for 2012 as no test strains for this year were available, so a rise could not be calculated); solid points show attack rate based on four-fold rise.

## Discussion

Our analysis shows that detailed mechanistic insights can be gained from longitudinal data by jointly considering individual infection histories and antibody dynamics acting at multiple timescales. Building on previous analyses [12, 15, 22, 23], we estimated that non-primary influenza exposures generate a short-lived broad humoural response and a persistent narrow response, with each accumulating and degrading to different degrees over the course of a human lifetime. As well as quantifying processes that shape the antibody response against different influenza strains, our results suggest that accounting for such dynamics leads to improved estimation of population attack rates.

The short-lived broad response, which we estimated makes the largest contribution to titres following infection, is likely to influence selection pressure imposed on the virus as a result of population immunity; it has been suggested that such short-term nonspecific immunity could explain the constrained genetic diversity of circulating influenza viruses [4]. Our results would therefore have implications for use of serology to investigate the evolutionary dynamics of influenza, and hence identify potential vaccine strain candidates [22, 24, 25]. If a large proportion of a population had recently experienced infection, it is likely that the short term antibody response would protect these individuals against strains occupying a large region of antigenic space. However, these strains may become more transmissible as the short-term response wanes.

Additional insight into such antibody dynamics could be generated directly using modern methods of sorting and sequencing individual B cells [26]. During non-primary infections, existing memory B cells that are genetically diverged from germline B cells generated during prior infections are rapidly stimulated. These B cells may reach high peripheral frequencies rapidly but, on average, have lower avidity against the current strain than they would have had against that host’s previous infections [27]. However, there will be competing demands on these cell lines to produce antibodies and possibly to differentiate further to increase their avidity. In addition to these memory cells, there is also the potential for the stimulation of germline B cells which may take longer to achieve functional peripheral frequencies but have higher avidity [28]. When observed during early infection, these new lineages would be much more similar to germline B cells, and would form fewer phylogenetic clades per sorted cell than the rapid response. Later during infection, cells making up the persistent response would be at higher frequencies and be more differentiated, but still form only few clades. Antigenic seniority [15] may arise because novel lineages during later life infections have to compete with existing lineages for antigenic stimulation [29, 30]. After infection, the memory frequency of the B cells making up the broad response likely returns to their pre-infection levels and the new B cells establish new subordinate memory populations. The aggregate effect of these mechanisms over a lifetime is consistent conceptually with the results we have presented (Fig. S9).

The modelling approaches described here also could be employed in evaluating the effectiveness of different vaccination strategies, which depends on an ability to reliably infer population attack rates. Moreover, the broader concept of multiple timescales of antibody response would have potential implications for the design of innovative vaccines, such as highly-valent vaccines [11]. If broad responses have shorter durations than narrow responses, then the tradeoff between current vaccines and other proposed candidates may be time dependent. Participants in trials of novel influenza vaccines should therefore be followed up over multiple seasons so that the dynamics of their immune reponse to both vaccination and natural infection can be assessed. At best, such vaccination against influenza A/H3N2 may stimulate a similar response to natural infection. However, there is evidence that vaccine-mediated immunity wanes quickly [31], that vaccine effectiveness declines after multiple immunisations [32], and that broad response against a novel subtype fades after repeated vaccination [33]. With appropriate data on serology and vaccination history, the differences in dynamics between the two processes could be elucidated using the model structure we have presented.

As well as examining differences in vaccination-mediated immunity and antibody response following natural infection, future empirical studies could refine our estimate for waning of the short-term response by collecting serological samples at intervals of less than one year. Alternatively, or additionally, having information on timing of confirmed influenza infection between sample collections would make it possible to constrain possible infection events, and hence improve estimates of short-term dynamics. In our model, we also accounted for individual-level heterogeneity in titres by including normally distributed error in our observation model. Our results suggest that this error parameter is well-identified (Table S2), but it would be challenging to examine other potential heterogeneity in antibody responses – such as age-specific biases – in more detail with the data available without making strong assumptions about the nature of such heterogeneity.

Our inference approach could be used in future to guide the design of studies to infer key aspects of antibody dynamics or to estimate historical attack rates. Joint analysis of infection history and antibody dynamics could provide more accurate information about infection rates, particularly in the years preceding sample collection, and inform studies that rely on robust attack rate estimates. As a result, such methods could help ensure that serological studies to examine influenza immunity profiles have adequate statistical power to test hypotheses and identify key mechanistic processes. Our approach is also likely to be applicable to other cross-reactive pathogens, such as dengue fever and Zika viruses [34].

## Methods

### Serological data

We used two publicly available datasets in our analysis. In the southern China data, crosssectional serology was taken in 2009 from 151 participants in Guangdong province in southern China and tested using microneutralization assays against a panel of nine strains: six vaccine strains (A/Hong Kong/1/1968, A/Victoria/3/1975, A/Bangkok/1/1979, A/ Beijing/353/1989, A/Wuhan/359/1995, and A/Fujian/411/2002) and three strains that circulated in southern China in recent years preceding the study (A/Shantou/90/2003, A/Shantou/806/2005, and A/Shantou/904/2008) [18, 15]. The Vietnam data included longitudinal serology collected between 2007–2012 from 69 participants in Ha Nam [19], with sera tested using haemagglutination inhibition (HI) assays against a panel of up to 57 A/H3N2 strains isolated between 1968–2008 [11]. All of the Vietnam participants were unvaccinated against influenza, and 19% of the southern China participants reported prior influenza vaccination. In analysis of both datasets, we represented antibody responses by log-titre. For a titre dilution of 10 ≤ *D* ≤ 1280, log-titre was defined as log_2_(*D*/10)+1. The minimum detectable titre in both datasets was 10, so a dilution <10 was defined to have a log-titre of 0. The maximum observable titre in both datasets was 1280, which corresponded to a log-titre of 8. There were nine possible observable log-titres in our analysis, ranging from 0 to 8. The antigenic summary path used to represent strains in our analysis was generated by fitting a two-dimensional smoothing spline through 81 points representing the published estimated locations of strains in ‘antigenic space’ [11] (Fig. S1).The positions of strains in such a space depends on the distance between influenza antigens and reference antisera as measured by titre in an HI assay [17]. In the model, we assumed that strains circulating between 1968 and 2012 were uniformly distributed along this summary path.

### Model of expected titre given infection history

We expanded a previous modelling framework designed for cross-sectional data [12] to include short- and long-term dynamics. For an individual who had previously been infected with strains in the set *X*, the expected log-titre against strain *j* depended on five specific antibody processes:

1. Long-term boosting from infection with homologous strain. If an individual had been infected with only one strain, they would exhibit a fixed log-titre against that strain, controlled by a single parameter, *µ*_1_.
2. Antigenic seniority acting via suppression of subsequent responses as a result of prior immunity. The titre against a particular strain was scaled by a factor *s*(*X, m*) = max{0, 1 − *τ* (*N_m_* − 1)}, where *N_m_* is the number of the strain in the infection history (i.e. the first strain is 1, the second is 2 etc.) and |*X*| is the total number of infections, and *τ* was a parameter to be fitted.
3. Cross-reactivity from antigenically similar strains. As titres were on a log scale, we assumed the level of cross-reaction between a test strain *j* and infecting strain *m ∈ X* decreased linearly with antigenic distance. This was controlled by *d*_1_(*j, m*) = max{0,1 − *σ*_1_*δ_mj_*}, where *δ_mj_* was the two-dimensional Euclidean antigenic distance between strains *j* and *m* (Fig. S1), and *σ*_1_ was a parameter to be fitted.
4. Short-term boosting, which waned over time. For an infecting strain *m*, this process was controlled by *μ*_2_*w*(*m*) = *μ*_2_ max{0,1 − *ωt_m_*}, where *μ*_2_ was a boosting parameter and *ω* was a waning parameter to be fitted, and *t_m_* was the number of years since infection with strain *m*. We constrained *ω* ≤ 1 when fitting the model to ensure identifiability, as *ω* = 1 or *ω* > 1 implies that *w*(*m*) = 0 for all *t_m_* > 0.
5. Cross-reactivity for the short-term response. The level of cross-reaction between a test strain *j* and infecting strain *m* was given by *d*_2_(*j, m*) = max{0,1 − *σ*_2_*δ_mj_*}, where *δ_mj_* was the antigenic distance between strains *j* and *m*, and *σ*_2_ was a parameter to be fitted.

To combine the five processes in the model, we assumed that the expected log-titre individual *i* had against a strain *j* was a linear combination of the responses from each prior infection:

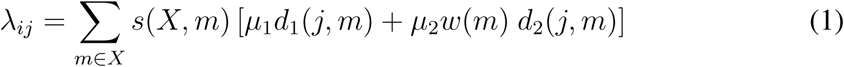

Depending on parameter values, our model could incorporate several specific mechanistic features, including: long-term response only (*μ*_2_ = 0); waning response only (*μ*_1_ = 0); or long-term/short-term boosting independent of a cross-reactive memory response (*σ*_1_, *σ*_2_ = 0).

### Observation model and likelihood function

For an individual *i* who was infected with strains in the set *X*, we assumed their true titre against strain *j* titre followed a normal distribution with mean *λ_ij_*, standard deviation *ε*, and cumulative distribution function *f* (*x*). The observed distribution of titres was censored to account for integer valued cutoffs. The likelihood of observing titre *k* ∈ {0, …, 8} given history *X* and parameter set *θ* was therefore as follows:

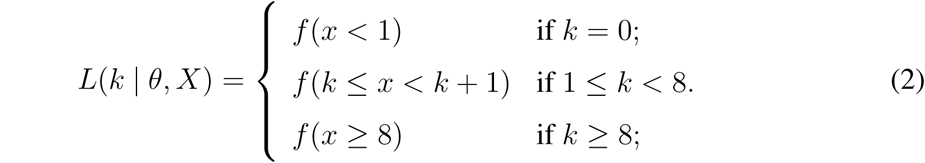

### Parameter estimation

We fitted the model to serological data using Markov chain Monte Carlo (MCMC). Using the likelihood function in Equation 2, we jointly estimated *θ* across all individuals and estimated *X* for each individual via a Metropolis-Hastings algorithm. If individual sera were collected in more than one year, parameters were jointly estimated across all test years. We used a data augmentation approach to estimate individual infection histories. Every second iteration, we resampled model parameters, which were shared across all individuals, and performed a single Metropolis-Hastings acceptance step. On the other iterations we resampled infection histories for a randomly selected 50% of individuals. These histories were independent across individuals, so we performed a Metropolis-Hastings acceptance step for each individual separately. Correlation plots indicated that all parameters in the full model were identifiable (Fig. S10). To ensure the Markov chain was irreducible, resampling at each step involved one of the following: addition of infection in some year; removal of infection in some year; moving an infection from some year to another [35]. We also used adaptive MCMC to improve the efficiency of mixing: at each iteration, we adjusted the magnitude of the covariance matrix used to resample *θ* to obtain an acceptance rate of 0.234 [36]. As we had data on participants individual ages in the southern China data, we constrained potential infections in the model to years in which participants would have been alive. The model was implemented in R version 3.3.1 and C, and used the Rcpp and doMC packages. Source code and data are available at: https://github.com/adamkucharski/flu-model/

### Simulation study

In our simulation study, we first generated simulated influenza attack rates between 1969– 2012 using a lognormal distribution with mean 0.15 and standard deviation 0.5. For 1968, we used a lognormal distribution with mean 0.5, to reflect higher incidence in the pandemic year [37]. Using these simulated attack rates, we generated individual infection histories for 69 participants using a binomial distribution, then generated observed individual level titres against the same strains as in the Vietnam dataset using our titre model. As in the real data, simulated samples were tested each year between 2007–2012. We assumed *μ*_1_ = *μ*_2_ = 2, *τ* = 0.05, *ω* = 1, *σ*_1_ = 0.3, *σ*_2_ = 0.1 and *ε* = 1 in simulations. For Fig. 3B inset, we simulated 10 independent sets of observed titres, then inferred the proportion of the population infected in the four years between 2008–2011 inclusive. The resulting distribution of model residuals (i.e. estimated minus actual simulated value) for these 40 data points were plotted as kernel density plots.

### Epidemiological data

Reported influenza A/H3N2 activity in Vietnam was obtained from the WHO FluNet database [38] (Fig. S11). We aggregated reports into temporal windows based on dates of serological sample collection [11], and used the cumulative number of isolates in each period to compare observed activity with model estimates. To calculate attack rates from the model outputs, we scaled the posterior distribution of total number of infections across all participants for each year between 1968–2012 by the proportion of participants who were alive in that year, which we calculated based on the age distribution of participants. This produced the estimates in Figs. 3C–D. A.J.K. was supported by the Medical Research Council (fellowship MR/K021524/1) and Sir Henry Dale Fellowship jointly funded by the Wellcome Trust and the Royal Society (Grant Number 206250/Z/17/Z). S.R was supported by the Medical Research Council (UK, Project MR/J008761/1), Wellcome Trust (093488/Z/10/Z, 200861/Z/16/Z, 200187/Z/15/Z), National Institute for General Medical Sciences (MIDAS U01 GM110721-01), National Institute for Health Research (UK, for Health Protection Research Unit funding). We would also like to thank Hua-Xin Liao for helpful discussions.

